# Contributions of temporal and spatial masking signals in perception of sequential visual events

**DOI:** 10.1101/2024.12.20.629621

**Authors:** Antimo Buonocore, Maria Cuomo, Martina Maresca, Alessio Fracasso

## Abstract

Accurate perception of time and space is essential for moment-to-moment interactions with our surroundings. This process requires flexibility, as it integrates information from our actions and the external context. Probing the visual system during the updating process reveals spatiotemporal distortions, where sequential stimuli appear closer in time and space than they are. These effects occur perisaccadically or when a visual mask follows the stimuli. The study investigated whether non-overlapping visual masks could influence temporal inversion judgments (TOJs), suggesting that a temporal signal might act as an anchor during updating. In Experiment 1, participants judged the temporal order of two stimuli under three conditions: no mask, a full-field mask, or a partial mask avoiding stimuli’s locations. Compared to no mask, both masks triggered TOJs when presented within 30 milliseconds of the second stimulus. In a control experiment, delaying mask onset by 30 milliseconds eliminated the inversion effect. In Experiment 2, TOJs were observed for both ipsilateral and contralateral masks, suggesting that long range inhibitory signals might also contribute to the effect. Together, these findings indicate that temporal inversions can occur with non-overlapping stimuli masks configuration, pointing to a non-spatial signal related to mask timing as the underlying mechanism.

## Introduction

Human perception is grounded in the assumption of continuity and sequential organization across both spatial and temporal dimensions of sensory input (He, Richter, Wang, & Lange, 2022; Wurtz, 2008). For example, the ability to intercept a moving object depends on motion extrapolation mechanisms (Nijhawan, 1994), that predict its spatiotemporal trajectory, thereby enabling precise perceptual and motor coordination (Kerzel & Gegenfurtner, 2003). Disruptions in these mechanisms —particularly in processing spatial or temporal order, can severely impair perceptual accuracy and goal-directed behavior.

However, a large body of research demonstrates systematic distortions in the perceived location and timing of stimuli presented in quick succession (Honda, 1991; Matin & Pearce, 1965). These distortions are particularly pronounced during the perisaccadic interval, the brief temporal window surrounding saccadic eye movements (Schlag & Schlag-Rey, 2002). Stimuli presented during this interval often appear shifted toward the saccade endpoint, a phenomenon known as perisaccadic compression (Bischof & Kramer, 1968; Buonocore & Melcher, 2015; Lappe, Awater, & Krekelberg, 2000; Morrone, Ross, & Burr, 1997; Ross, Morrone, & Burr, 1997). In the temporal domain, when two stimuli are presented sequentially, one before and one after a saccade, their temporal separation is frequently underestimated (Binda, Cicchini, Burr, & Morrone, 2009; Morrone, Ross, & Burr, 2005). In some cases, this temporal compression can even result in a reversal of perceived order, with the later stimulus erroneously perceived as occurring earlier (Morrone et al., 2005).

These perceptual errors offer critical insights into the brain’s functioning, suggesting that the neural mechanisms underlying spatial and temporal integration are transiently altered. At the neurophysiological level, theories propose that the brain employs extraretinal signals to compensate for retinal shifts caused by saccades. These signals include an efference copy (Holst & Mittelstaedt, 1950) or corollary discharge (Sperry, 1950) of the saccadic command, which counteracts the effects of retinal image translation. A key explanation for spatial compression involves receptive field shifts in anticipation of eye movements, driven by corollary discharge (Melcher & Colby, 2008; Wurtz, 2008). As the eyes prepare to move, the brain dynamically adjusts its spatial representation, leading to temporary distortions in perceived object positions that can be probed during the updating interval. Regions such as V2, V3 (Nakamura & Colby, 2002), V4 (Tolias et al., 2001), the lateral intraparietal area (LIP) (Duhamel, Colby, & Goldberg, 1992), the frontal eye fields (FEF) (Joiner, Fitzgibbon, & Wurtz, 2010; Shin & Sommer, 2012; Umeno & Goldberg, 1997), and the superior colliculus (SC) (Walker, Fitzgibbon, & Goldberg, 1995) are involved in these predictive adjustments, modulating sensory input to minimize the perception of retinal displacement. An alternative explanation points to mislocalization errors as a result of receptive fields shifting to the saccade target location, rather than their future location. Zirnsak and colleagues (2014) showed that FEF receptive fields are preferentially activated near the saccade target already during the presaccadic period, causing a population response shift toward the saccade endpoint, where the perceptual mislocalization is also recorded (Zirnsak, Steinmetz, Noudoost, Xu, & Moore, 2014).

Temporal distortions, on the other hand, are more complex and are often attributed to saccadic suppression and visual masking effects. When stimuli are presented across a saccade, one before and one shortly afterward, the brain attempts to integrate pre- and post-saccadic visual information, but this integration is often imperfect. Saccadic suppression (Bremmer, Kubischik, Hoffmann, & Krekelberg, 2009; Thiele, Henning, Kubischik, & Hoffmann, 2002) can delay encoding of the first stimulus, causing the timing to be misjudged. This misjudgment can manifest as temporal compression, where the first stimulus appears closer in time to the second, or even as a reversal of perceived order, with the second stimulus erroneously perceived as occurring first. Another explanation involves overlapping probability distributions, where the brain’s estimation of time relies on probabilistic models of event timing. When these distributions overlap, temporal intervals may be perceived as condensed, resulting in distortions in perceived order and duration (Binda et al., 2009).

Not all spatial-temporal distortions are directly tied to eye movements. Similar effects have been observed during stable fixation, particularly when the visual scene is disrupted. For example, masks mimicking saccadic retinal shifts (Mackay, 1970; O’Regan, 1984; Ostendorf, Fischer, Gaymard, & Ploner, 2006), flicker stimuli (Terao, Watanabe, Yagi, & Nishida, 2008), or brief visual masks (Buonocore & Melcher, 2015; Chota et al., 2020; Zimmermann, Born, Fink, & Cavanagh, 2014) can elicit spatial and temporal compression effects akin to those seen during saccades. These findings suggest that visual masking may act as a functional analog of retinal shift signals, effectively triggering the same neural mechanisms responsible for perisaccadic distortions and being responsible for visual continuity (Zimmermann et al., 2014).

Currently, it remains unclear which specific characteristics of the mask underlie the observed effects. One hypothesis is that the mask’s visual properties mimic the effects of saccades by disrupting spatial and temporal continuity. Alternatively, we suggest that the onset of the mask may provide a timing signal that serves as an anchor for organizing the temporal sequence of stimuli. When combined with masking-induced interruptions in vision, this timing signal could account for the full range of spatiotemporal distortions described in the literature. This hypothesis aligns well with findings on mislocalization under a saccadic inhibition paradigm (Buonocore & Melcher, 2015). In those experiments, the authors demonstrated that delaying a saccade during its programming phase by means of saccadic inhibition (Buonocore & McIntosh, 2008; Edelman & Xu, 2009; Reingold & Stampe, 2002) shifted the mislocalization error backward in time, suggesting a link between saccadic programming and perception. This implies that saccade preparation may be the key driver of perceptual changes, rather than solely the visual masking component of the saccade. Similarly, in the present design, the mask might provide a timing signal that is independent of its visual characteristics.

To disentangle these hypotheses, we conducted experiments in which participants made temporal order judgments for two visual stimuli presented sequentially under varying masking conditions. The masks differed in their spatial overlap with the stimuli. If the inversion effect requires complete mask-stimulus overlap, this would support a purely visual explanation. However, if partial masks induce TOJ reversals, it would suggest that a temporal signal also plays a role. In the following experiments, we systematically examine the relationship between visual masking and TOJ reversals to clarify these mechanisms.

## Methods

### Experiment 1 – Partial masking

In Experiment 1, we investigated whether TOJ about the presentation order of two visual stimuli —a test and a probe, could be reversed using a visual mask that did not occlude the locations of the perceptual stimuli. Whitin this context, the mask is causing only a weak visual disruption but it still provides a clear signal about its onset. We hypothesized that this mask-related temporal signal might serve as an anchor to reorganize the order of the presented stimuli, similarly to what it has been observed by shifting saccades in time during perisaccadic mislocalization (Buonocore & Melcher, 2015). Furthermore, the reversal of TOJ might also be strengthened by a long-range spatial inhibitory mechanism between the mask and the test/probe locations. To test this hypothesis, we designed a novel experiment that involved both a full mask and a partial mask, presented at different intervals after the probe stimulus. In the full mask condition, we aimed to replicate the findings of Chota and colleagues (2020), which demonstrated that TOJ reversal increased when a full masking stimulus was presented within 30 ms of the probe stimulus (Chota et al., 2020). To test our specific hypothesis, we introduced a new masking condition in which the mask covered only the top and bottom thirds of the screen, leaving the discrimination stimuli fully visible. If TOJ reversal occurs under this condition, it would suggest that factors other than visual occlusion —such as the temporal signals generated by the mask, may play a role in inducing the reversal.

### Participants

In Experiment 1, fourteen participants (10 females; mean age = 25.86 years, SD = 5.72) were recruited to participate in the study. All participants were free from neurological, psychiatric and visual impairments, and were naive to the purposes of the experiment. Participants were students from the university and had normal or corrected to normal vision. Written informed consent was obtained, in accordance with the 1964 Declaration of Helsinki. Ethical approval was granted by the local ethics committee at the college of Medical, Veterinary and Life Sciences, University of Glasgow.

### Sample size consideration

The experiments reported in the paper are based on the paradigm described by Chota and colleagues (2020), which demonstrates a strong effect of TOJ reversals induced by masking stimuli closely following the last stimulus in the sequence. For this reason, we used a sample size (N=14) comparable to that in the referenced article (N=14).

### Apparatus

Participants were positioned in a chin-and-forehead rest apparatus to ensure head stability. Responses were recorded using a standard keyboard. Visual stimuli were presented on a 27-inch monitor with a resolution of 1920 by 1080 pixels and a refresh rate of 144 Hz. The participants’ eyes were aligned with the center of the display at a distance of about 50 cm. The experiment was implemented in Matlab (R2021a, The MathWorks, Inc., Natick, MA), utilizing the Psychtoolbox (Brainard, 1997; Pelli, 1997).

### Stimuli

The stimuli in Experiment 1 included a blue dot (0.22 degrees of visual angle, DVA) serving as a fixation point positioned at the center of the screen, two rectangles (0.74 by 4.61 DVA) —one red (RGB: [255, 0, 0]) and one yellow (RGB: [255, 234, 0]) —used for temporal discrimination judgments, and various types of masks. All stimuli were presented against a mid-gray background (RGB: [128, 128, 128]). Following the terminology of Chota and colleagues (2020), the red rectangle, referred to as the "target," was located 12.5 DVA to the right of the fixation point at mid-screen height. The yellow rectangle, referred to as the "probe," was positioned 2.6 DVA to the right of the target. Both the target and probe were displayed for 21 milliseconds, and the mask stimulus appeared 160 milliseconds after the first stimulus and lasted for 56 milliseconds. Stimulus timing was calculated based on a 144 Hz screen refresh rate.

In Experiment 1, both the timing of the probe relative to the target stimulus and the type of mask were manipulated. The probe stimulus appeared at various stimulus onset asynchronies (SOAs) ranging from -56 milliseconds to 132 milliseconds (-56, -35, 56, 111, 132 ms). For conditions where the probe appeared after the target, the mask followed the probe by 104, 49, and 28 milliseconds, respectively, in this way we could test whether temporal proximity between mask and the probe would affect TOJ responses, while keeping the presentation of the target unaltered. The experiment involved three masking conditions: a control condition with no masking stimulus, a full mask condition in which the mask covered the entire screen, and a partial mask condition where the mask consisted of two rectangles covering only the top and bottom thirds of the screen, with heights equivalent to one-third of the screen height and a width equal to the screen width, leaving the central area uncovered. In both the full mask and partial mask conditions, the masking stimulus was composed of pixels with random luminance values ranging from RGB [0, 0, 0] to RGB [255, 255, 255]. The experiment included 15 conditions, resulting from the factorial combination of mask type (no mask, full mask, partial mask) and probe SOA (-56, -35, 56, 111, 132 ms), which were presented in random order. Each condition was repeated 30 times, with participants completing 10 blocks of 45 trials each, for a total of 450 trials per participant. The experimental session lasted approximately 30 minutes, including breaks.

### Procedure

In each trial, participants were instructed to maintain fixation on the central dot for the entire duration of the trial (Figure 1A). After a random interval of 1000 to 1500 milliseconds following the appearance of the fixation dot, either the target or the probe stimulus was presented on the right side of the screen. In some trials, either a full mask or a partial mask was displayed 160 milliseconds after the first stimulus. Following the stimulus presentation, a blank screen appeared with a question prompting the participant to indicate which stimulus was presented first. Participants reported whether the red rectangle (the target) or the yellow rectangle (the probe) appeared first by pressing the corresponding keys on the keyboard, with "a" indicating that the red rectangle appeared first and "l" indicating that the yellow rectangle appeared first. There was no time limit for giving a response, and the next trial began immediately after a response was provided. Participants were allowed to take a short break after each block of trials and could start the next block by pressing any key on the keyboard. At the end of the session, all participants were debriefed regarding the aim of the research.

**Figure 1.**
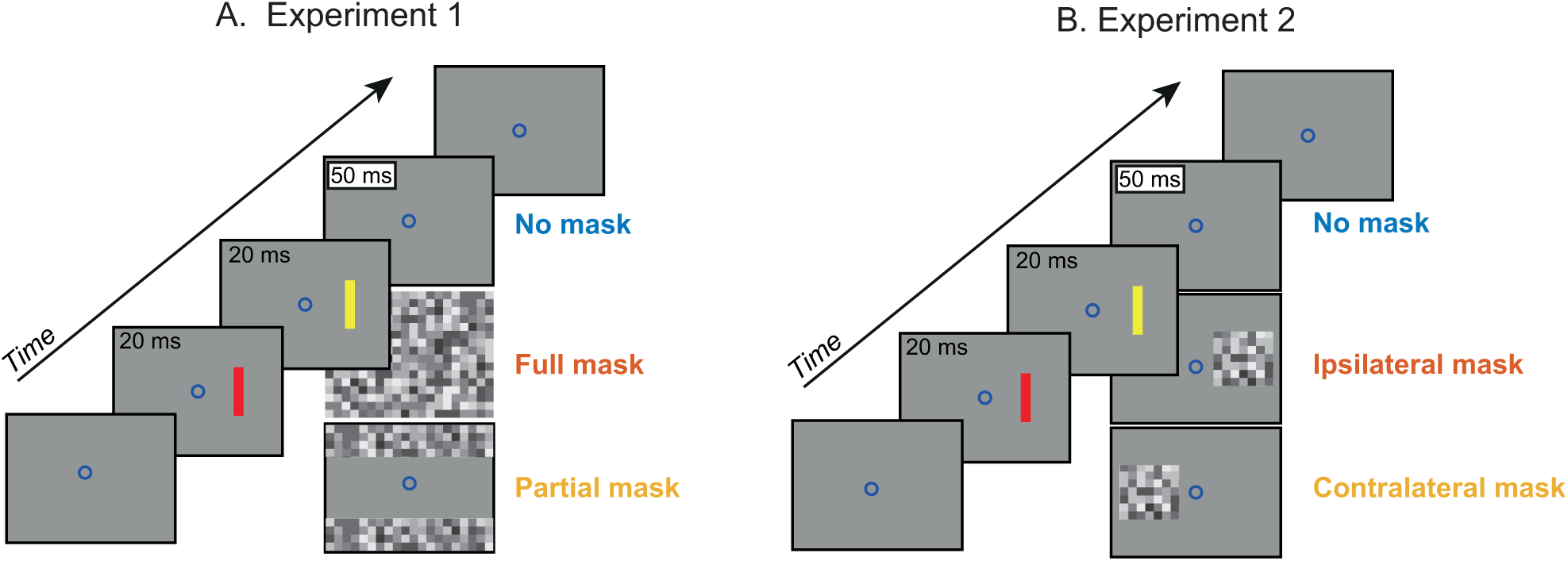
Stimulus paradigms. (A) In Experiment 1, participants maintained fixation on a blue dot throughout the trial. After a variable delay from trial start (1000–1500 ms), either a red target or a yellow probe was presented for 20 ms. The probe could precede the target onset by -56 ms or -35 ms or follow it by 56 ms, 111 ms, or 132 ms. A full-field pixelated mask was displayed for 56 ms at 160 ms after the onset of the first stimulus. The experimental design included three conditions: no mask (blue), a full-field mask (red), and a partial-field mask (yellow). (B) In Experiment 2, the sequence of events was identical to Experiment 1, but the mask was either ipsilateral (red) or contralateral (yellow). A no-mask condition (blue) served as the baseline.

## Data analysis

All data analyses were conducted using MATLAB (MathWorks). To compute group averages, we first calculated the probability of responding "probe first" for each participant (N = 15) and each experimental condition (15 conditions: mask type by probe SOA). We then computed the standard error of the mean across the sample for each condition. To investigate how the perception of the probe ("probe first" coded as 0 or 1) was influenced by mask type (categorical predictor: no mask, full mask, partial mask), probe SOA (continuous predictor: -56, -35, 56, 111, 132 ms), and their interaction, we fit a mixed-effects logistic regression (GLME). The model included participants as a random effect to account for individual variability. The structure of the model is summarized in Wilkinson notation (Wilkinson & Rogers, 1973) as follows:

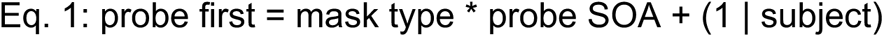

To follow up on significant interactions, we calculated the increase in the probe first probability by subtracting the response probability in each masking condition from the corresponding probability in the no-mask condition. Positive values indicated an increase in the reversal of temporal order judgments (TOJ). We assessed significant differences by performing t-tests against zero for the positive probe SOA. All critical p-values were set to 0.05.

## Results

### Partial Masking Induces TOJ Reversals

We first investigated whether the perception of probe was influenced by the presence of a mask and whether a partial mask could induce TOJ reversals. To address this, we fit a mixed-effects logistic regression analysis using "probe first" (0, 1) with mask type (no mask, full mask, partial mask), probe SOA (-56, -35, 56, 111, 132 ms), and their interaction as predictors. The model is summarized in Equation 1.

The analysis revealed that probability in detecting whether the probe appeared first was significantly modulated by probe SOA. The GLME analysis (see Table 1) identified a significant main effect of probe SOA (β = -0.023, 95% CI [-0.0245, -0.021], t = -26.14, p < 0.001). This result indicates that probes presented before the test stimulus were correctly identified with a probability exceeding 80%, which sharply declined to approximately below 20% when the probe was presented after the test. These findings suggest that the task was not at ceiling and participants wrongly reported the order of target-probe about 15% of the times, even in the absence of a mask, and for SOA over 100ms.

**Table 1.** Experiment 1 GLME parameters.

| Name | Beta | S.E. | t | DF | p | 95% CI<br>lower | 95% CI<br>upper |
| --- | --- | --- | --- | --- | --- | --- | --- |
| Intercept | 0.557 | 0.182 | 3.066 | 6214 | 0.002 | 0.201 | 0.914 |
| **SOA** | **-0.023** | **0.001** | **-26.140** | **6214** | **<0.001** | **-0.025** | **-0.021** |
| Mask (full) | 0.172 | 0.092 | 1.870 | 6214 | 0.061 | -0.008 | 0.353 |
| Mask (partial) | 0.108 | 0.091 | 1.179 | 6214 | 0.238 | -0.071 | 0.287 |
| **SOA : Mask (full)** | **0.004** | **0.001** | **3.231** | **6214** | **0.001** | **0.001** | **0.006** |
| **SOA : Mask (partial)** | **0.002** | **0.001** | **2.018** | **6214** | **0.044** | **0.000** | **0.005** |

More importantly, we found a significant two-way interaction between probe SOA and mask type. Specifically, both the full mask (β = 0.004, 95% CI [0.0014, 0.006], t = 3.2314, p = 0.006) and the partial mask (β = 0.002, 95% CI [0.0001, 0.0047], t = 2.018, p = 0.044) significantly interacted with probe SOA. These results suggest that the presence of a mask increased the likelihood of TOJ reversals. Figure 2A shows the average probe-first frequency scores for the interaction between probe SOA and mask type, emphasizing an increase in the probability of reporting probe first for both the full mask (red) and the partial mask (yellow).

**Figure 2.**
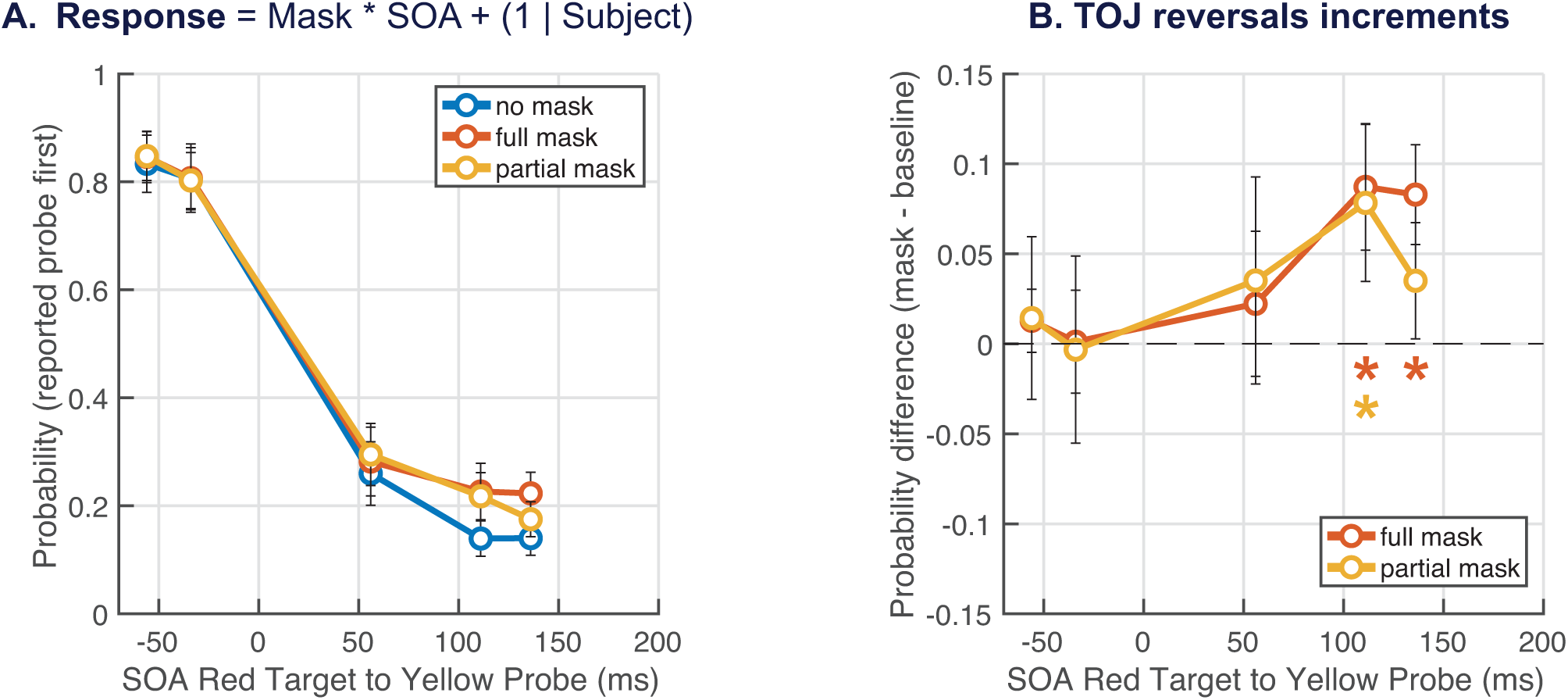
Mean results from Experiment 1 (N = 14). (A) Average frequency for “probe first” presented across all conditions. The data are split by the mask type: no mask condition (blue), full mask (red), and partial mask (yellow). (B) Difference between full mask (red), and partial mask (yellow) condition against the no mask one.

A positive value indicates an increased proportion of TOJ reversals. All error bars represent one SEM. Asterisks represents significant pairwise t-test with an alpha level lower than 0.05.

We extended the analysis by calculating the increase in the probability of reporting probe first by subtracting the probabilities in each masking condition from those in the no-mask condition. In this analysis, values greater than zero indicated an increase TOJ reversal, which was tested using paired t-tests for the critical levels of probe SOA, when it was following the test stimulus (Figure 2B). The results showed that the increase in TOJ reversal was significantly greater than zero at 49 ms (t = 2.446, df = 13, 95% CI [0.0095, 0.1525], p = 0.029) and 26 ms (t = 3.277, df = 13, 95% CI [0.0287, 0.1396], p = 0.006) before the full mask. Similarly, the partial mask condition showed a significant increase at 49 ms (t = 3.378, df = 13, 95% CI [0.0245, 0.1114], p = 0.005).

These findings suggest that visually masking the stimulus was not the primary factor inducing TOJ reversal. Instead, the results point to a temporal component, likely generated by the mask onset, that acts as an anchor signal for reorganizing the visuo-temporal perception of the stimuli. Furthermore, the data indicate that a long-range visual signal may still contribute to the masking effects observed on the screen (Fabius, Fracasso, & Stigchel, 2016).

### Experiment 1 – Control

In a control for Experiment 1, we placed the mask further away from the probe stimuli. Moreover, to strengthen the possible TOJ reversal effect, we moved the fixation on the left side of the screen, so that participants were potentially ready to prepare eye movements even if they were not required to do so.

## Methods

### Participants

In the control experiment, seven participants (6 females; mean age = 24.43 years, SD = 2.15). All participants reported normal or corrected to normal vision.

### Stimuli and procedure

The stimuli and procedure were identical Experiment 1, except for the following changes. In Experiment 1 – control, the full-field mask or a partial-field mask were presented 180 ms after the onset of the first stimulus in the sequence, i.e. with 30 ms delay compared to Experiment 1. This means that the minimum time between probe onset and mask was 46 ms, corresponding at probe SOA of 134 ms. Moreover, the fixation point was a black cross, situated toward the left of the screen (7.2 of visual angle from midline), at mid height, and was present continuously. The eccentric position of the fixation cross served to simulate conditions for an experiment with saccades toward the targets, albeit participants were strictly required to not make any eye movement (Chota et al., 2020).

## Results

### Late masks do not cause TOJ reversals

In the control experiment, the only significant result was the effect of probe SOA (β = -0.020, 95% CI [-0.0219 -0.0175], t = -17.43, p < 0.001), suggesting that participants were correctly performing the task (see Table 2). Similarly to Experiment 1, all conditions never reached ceil or floor levels, suggesting that there was always a proportion of TOJ reversals.

**Table 2.**
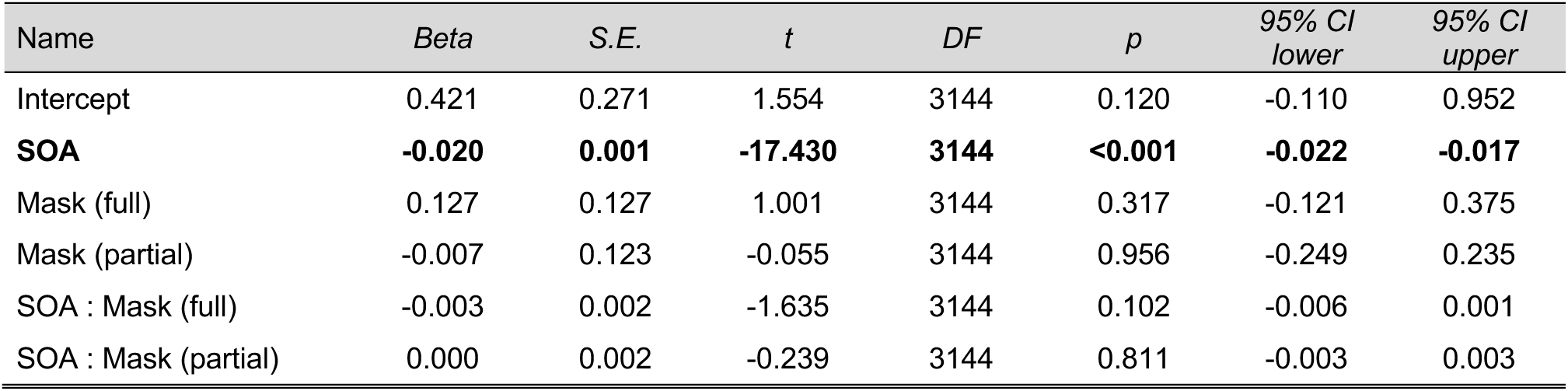
Experiment 1 - control GLME parameters.

The lack of any main effect of mask and interactions supports the idea that the mask needs to be closely following in time the onset of the probe to induce the TOJ reversal (Figure 3A and B).

**Figure 3.**
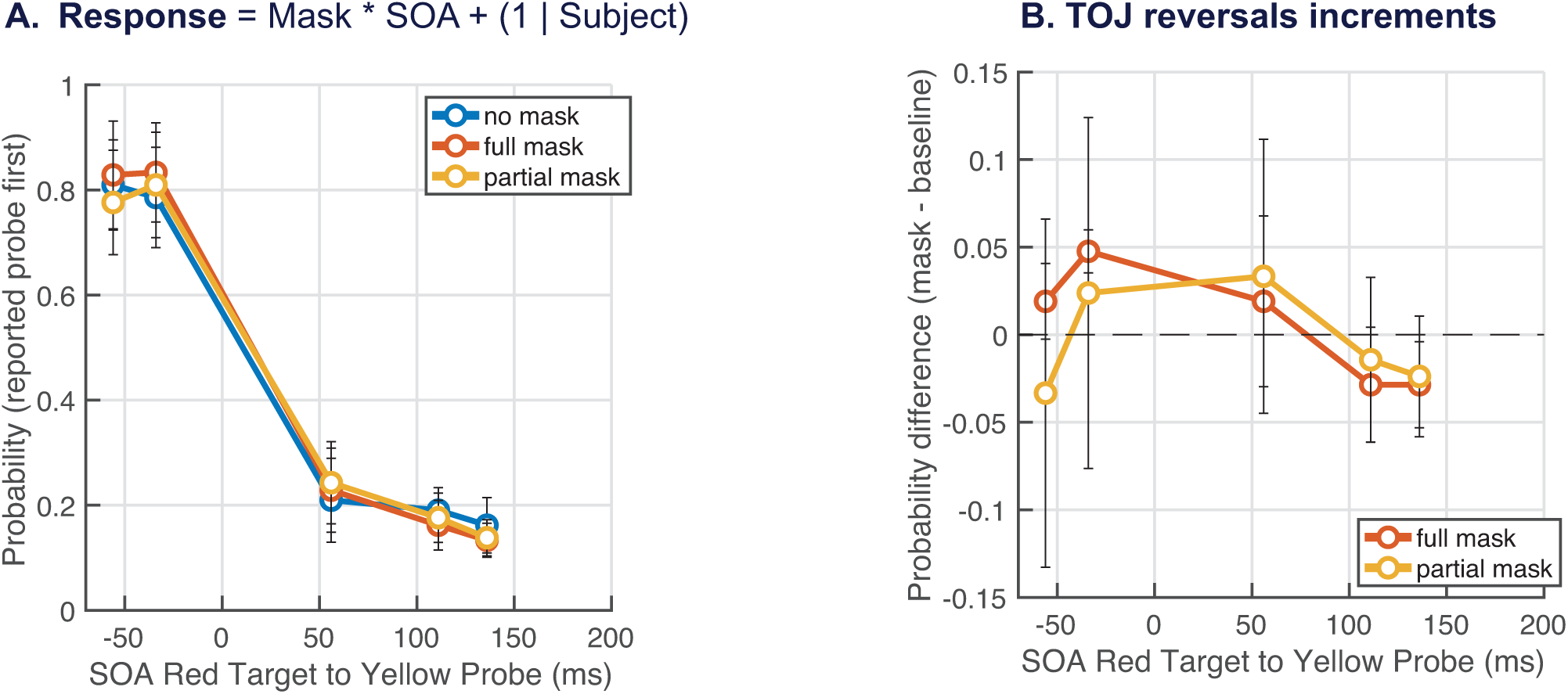
Mean results from Experiment 1 – control (N = 7). Same convention as Experiment 1 but now the mask is presented 180 ms after first stimulus in the sequence. (A) Average frequency for “probe first” (B) Difference between masks conditions and the no mask one. Differently from Experiment 1, no significant increase in TOJ was observed. All error bars represent one SEM.

### Experiment 2 – Lateralized masking

In Experiment 2, we further tested the idea of partial masking by moving the mask in one experimental condition in the contralateral hemifield. In this case, we reduced the potential impact of within-field, long range inhibitory spatial effect (Fabius, Fracasso, Nijboer, & Stigchel, 2019; Fabius et al., 2016; A Fracasso, Caramazza, & Melcher, 2010) . In the case of TOJ reversals for the contralateral stimuli, we hypothesize a strong contribution of a temporal signal rather than a spatial one.

## Methods

### Participants

In the second experiment fifteen participants (10 females; mean age = 22.9 years, SD = 2.8) participated in Experiment 2. All participants reported normal or corrected to normal vision.

### Stimuli and procedure

The stimuli and procedure were identical to Experiment 1, except for the following changes. In Experiment 2, instead of a full-field mask or a partial-field mask, a square mask is presented either in the ipsilateral visual field, covering only the two rectangular stimuli, or in the contralateral hemifield relative to the two stimuli. The mask was a square with dimensions of 10 DVA, with square pixels measuring 0.5° each of which assumed a random luminance value (RGB ranging from 0 to 255).

## Results

### A temporal signal drives TOJ reversals

We run the same analysis as Experiment 1, testing the effect of mask type and probe SOA on a GLME (see Equation 1). As experiment 1 and control, we confirmed a main effect of probe SOA (β = -0.018, 95% CI -0.0120 -0.0167], t = -24.775, p < 0.001). But more importantly, we also replicated the significant interaction between mask type and probe SOA (see Table 3).

**Table 3.**
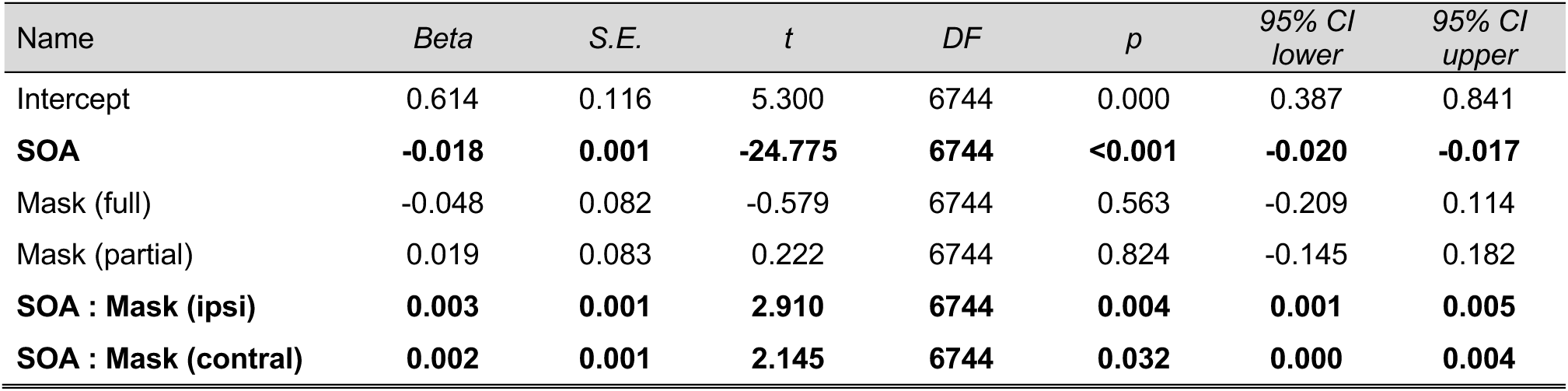
Experiment 2 GLME parameters.

| Name | Beta | S.E. | t | DF | p | 95% CI lower | 95% CI upper |
| --- | --- | --- | --- | --- | --- | --- | --- |
| Intercept | 0.614 | 0.116 | 5.300 | 6744 | 0.000 | 0.387 | 0.841 |
| **SOA** | **-0.018** | **0.001** | **-24.775** | **6744** | **<0.001** | **-0.020** | **-0.017** |
| Mask (full) | -0.048 | 0.082 | -0.579 | 6744 | 0.563 | -0.209 | 0.114 |
| Mask (partial) | 0.019 | 0.083 | 0.222 | 6744 | 0.824 | -0.145 | 0.182 |
| **SOA : Mask (ipsi)** | **0.003** | **0.001** | **2.910** | **6744** | **0.004** | **0.001** | **0.005** |
| **SOA : Mask (contral)** | **0.002** | **0.001** | **2.145** | **6744** | **0.032** | **0.000** | **0.004** |

We further investigated the two interactions by calculating the difference between each mask type and the no-mask condition at each probe SOA (Figure 4B). Consistent with the findings from Experiment 1, we observed that both mask types induced a significant increase in TOJ reversals shortly before the mask onset. Specifically, in the ipsilateral condition, the increase was significant 49 ms before the mask onset (t = 2.402, df = 14, 95% CI [0.0086, 0.1514], p = 0.030). In the contralateral mask condition, a significant increase in TOJ reversals was observed 28 ms before the mask onset (t = 2.537, df = 14, 95% CI [0.0096, 0.1148], p = 0.024).

**Figure 4.**
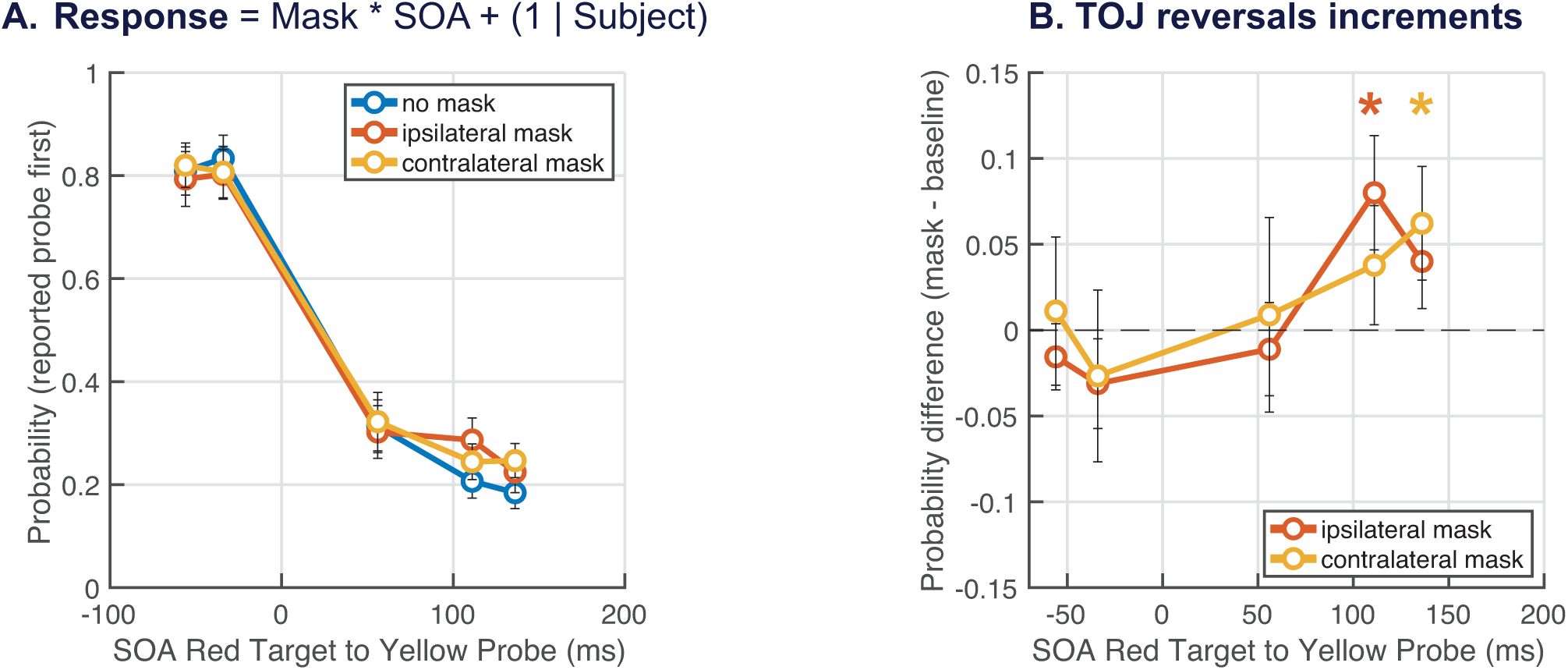
Mean results from Experiment 2 (N = 15). Same convention as Experiment 1 but now the mas can be presented ipsilaterally (red) or contralaterally (yellow). (A) Average frequency for “probe first”. (B) Difference between mask conditions against the no mask one. Similarly to Experient 1, the last two probe SOA indicate an increase in TOJ reversals. All error bars represent one SEM. Asterisks represents significant pairwise t-test with an alpha level lower than 0.05.

The results of both Experiment 1 and Experiment 2 indicate that the mask reliably induces TOJ reversals. More importantly, these findings demonstrate that this effect is not primarily driven by the mask spatially occluding the stimuli. Both the partial mask and contralateral mask conditions reliably induced TOJ reversals. These results suggest that a timing signal associated with the onset of the mask is the primary driver of the effect. This timing signal may operate in conjunction with long-range inhibitory interactions on the spatial map, which contribute to—but are not solely responsible for—the perceptual ordering of the stimuli.

## Discussion

This study demonstrated the illusory reversal of temporal order judgments for pairs of stimuli briefly presented before a visual mask (Figs. 2, 4). Specifically, when a visual mask closely followed a probe in time, the probe was perceived as occurring earlier than a test stimulus presented up to 130 ms prior. This finding aligns with earlier observations of spatial distortions reported in saccadic paradigms (Binda et al., 2009; Buonocore & Melcher, 2015; Kresevic, Marinovic, Johnston, & Arnold, 2016; Morrone et al., 2005), where eye movements caused a perceptual compression of time and space for stimuli presented before and after a saccade. Furthermore, it replicates findings from visual masking studies conducted during fixation (i.e., in the absence of eye movements), which revealed similar spatiotemporal distortions (Zimmermann et al., 2014), suggesting that these effects are not saccade- specific but instead reflect a more general mechanism of disruption-induced perceptual reorganization.

Focusing on previously documented TOJ reversals induced by visual masks (Chota et al., 2020), our results extend these findings to conditions where the masking stimuli do not spatially overlap with the test or probe stimuli. In Experiment 1, we demonstrated TOJ reversal using a mask with a central cutout to avoid spatial overlap (Fig. 1A). In Experiment 2, we further extended these findings to masks positioned in the opposite hemifield from the stimuli (Fig. 1B). Together, these results suggest that the reorganization of temporal order does not require direct spatial interactions between the mask and the stimuli, as previously proposed in studies examining spatiotemporal distortions during saccades (Morrone et al., 2005) and visual masking (Chota et al., 2020; Zimmermann et al., 2014). Instead, any temporal signal generated by a visual stimulus (in this case, the mask) which is in close temporal proximity to the probe may be sufficient to alter the perceived order of briefly presented visual events.

Zimmermann and colleagues (2014) interpreted results from visual masking paradigms as evidence that visual disruptions, such as saccades or masking, trigger a process of serially updating the scene representation by linking pre- and post-disruption items. Disruptions occurring shortly after a probe may delay the target’s updating, effectively postponing its perception in the temporal sequence. The author suggested that this process likely relies on delayed feedback from higher-order areas (Binda et al., 2009; Enns, Lollo, Enns, & Lollo, 2000; Fahrenfort, Scholte, & Lamme, 2007). Our findings build on this idea by suggesting that the critical temporal disruption is not exclusively tied to visual masking processes. Instead, the temporal reordering of stimuli may result from any sufficiently salient visual event occurring after the test-probe sequence. These results also align with the eye movement literature, where Buonocore and Melcher (2015) demonstrated that for the spatial domain, perisaccadic mislocalization can shift in time based on the intention to move the eyes, rather than the movement itself. In their study, delays in saccade onset were introduced by means of irrelevant visual transients inducing saccadic inhibition (Buonocore & McIntosh, 2008; Edelman & Xu, 2009; Reingold & Stampe, 2002). Temporal reconstruction of the mislocalization curve for delayed saccades was shifted earlier, indicating that these errors were driven by the timing of saccadic preparation (motoric) (Buonocore & Hafed, 2023) rather than by the execution of the saccade itself. In In the present experiment as well as in Buonocore and Melcher (2015) the timing signal, either motoric or visual, is what matters in shaping perception.

One could ask to which extent the visual component of the mask is critical in the TOJ reversals. To address this point, a second mechanism has been proposed to account for TOJ, based on salience suppression (Kresevic et al., 2016). Within this framework, visual disruptions, including masking and saccades, can reduce the effective contrast or salience of stimuli, particularly those presented temporally close to the disruption. In TOJ tasks, lower-contrast stimuli are often perceived as occurring earlier than higher-contrast ones. To be consistent with this explanation, we propose that in our experiment’s salience suppression might be achieved by long-range spatial interactions (Spillmann & Werner, 1996) between the peripheral mask and the more parafoveal stimuli. These sort of lateral connections, typically found in visual areas with retinotopic maps such as V1 (Alessio Fracasso, Petridou, & Dumoulin, 2016; Macknik & Livingstone, 1998; Wandell, Dumoulin, & Brewer, 2007), may play a role by either introducing noise into the visual signal (Benzi, Sutera, & Vulpiani, 1981; Groen, Tang, Wenderoth, & Mattingley, 2018; O. Nakamura & Tateno, 2019; Simonotto et al., 1997) or via lateral inhibitory processes (Tzvetanov & Simon, 2006). Either way, the mask might work by reducing the probe salience through weak masking, and further affecting the perceived timing of stimuli in a similar manner as Kresevice and colleagues (2016) reported. However, salience suppression effects may only partially explain our data give the spatial configuration of the mask and test/probe stimuli. In Experiment 2 the contralateral mask is also inducing higher TOJ reversals, implying an interhemispheric masking effect. Rather, we suggest that the main driver of the illusion still relies more on the temporal signal generated by the onset of a masking stimulus which act as an anchor for reorganizing perceived events, while the contribution of the spatial signal is weaker.

The absence of TOJ reversals when the mask onset was delayed as in Experiment 1 – control, underscores the existence of a critical timing window, reinforcing the notion that temporal precision is essential for perceptual reorganization. Interestingly, previous models of temporal compression, such as those proposed by Binda and colleagues (2009), suggest that TOJ reversals result from overlapping probability distributions of target and probe timing. This model predicts reversals for very short intervals in the range of tenths of milliseconds, as greater overlap leads to stronger compression effects. However, it does not adequately explain our findings at longer intervals over 100 ms, where TOJ reversals still occurred, and for mask presented as far as 50 ms after the probe, in which TOJ disappears, suggesting that compression of time alone cannot account for these effects.

Finally, the results of our study can be interpreted within the framework of rise-to-threshold models of vision, which propose that the perception of visual events depends on the accumulation of sensory evidence until it reaches a threshold (Carpenter & Williams, 1995; Reddi & Carpenter, 2000). According to these models, each visual stimulus generates a neural signal that increases over time, with the rate and trajectory influenced by factors such as salience, contrast, and temporal proximity to other stimuli. In our experiments, the mask onset may disrupt this process by introducing a competing neural signal. This signal could either: (i) delay evidence accumulation for the test stimulus, postponing its rise to threshold and altering perceived temporal order; or (ii) introduce noise that allows the probe stimulus to reach threshold earlier than the test, despite being presented later (Benzi et al., 1981; Groen et al., 2018; O. Nakamura & Tateno, 2019; Simonotto et al., 1997). While in the case of TOJ the latter explanation might align with factors like salience suppression, the former is less plausible. A slight temporal shift in the mask’s onset, as in the control condition of Experiment 1, disrupts temporal order judgments across all conditions (full and partial). Thus, we suggest that the mask might accelerates the rise-to-threshold process by prioritizing the more salient stimulus. This view eliminates the need for probabilistic or serial processing accounts, emphasizing the importance of salience in visual perception.

In conclusion, this study highlights the importance of temporal signals in reorganizing perceived event order, even in the absence of spatial overlap between masking stimuli and visual events. These findings support the view that both temporal and spatial dynamics interact to shape perceptual outcomes. Future research should aim to uncover the neural circuits mediating these effects, and model them effectively, starting from first principles, bridging theoretical models with experimental data to refine our understanding of sensory integration in dynamic visual environments.

## Data availability statement

Original data will be made available on reasonable request.

